# Mn^2+^ coordinates Cap-0-RNA to align substrates for efficient 2′-*O*-methyl transfer by SARS-CoV-2 nsp16

**DOI:** 10.1101/2021.01.31.429023

**Authors:** George Minasov, Monica Rosas-Lemus, Ludmilla Shuvalova, Nicole L. Inniss, Joseph S. Brunzelle, Courtney M. Daczkowski, Paul Hoover, Andrew D. Mesecar, Karla J. F. Satchell

## Abstract

Capping viral messenger RNAs is essential for efficient translation and prevents their detection by host innate immune responses. For SARS-CoV-2, RNA capping includes 2′-*O*-methylation of the first ribonucleotide by methyltransferase nsp16 in complex with activator nsp10. The reaction requires substrates, a short RNA and SAM, and is catalyzed by divalent cations, with preference for Mn^2+^. Crystal structures of nsp16-nsp10 with capped RNAs revealed a critical role of metal ions in stabilizing interactions between ribonucleotides and nsp16, resulting in precise alignment of the substrates for methyl transfer. An aspartate residue that is highly conserved among coronaviruses alters the backbone conformation of the capped RNA in the binding groove. This aspartate is absent in mammalian methyltransferases and is a promising site for designing coronavirus-specific inhibitors.

---

The recently emerged human pathogenic SARS-CoV-2 is a positive-stranded RNA virus responsible for the on-going pandemic of highly transmissible fatal respiratory coronavirus infectious disease (COVID-19), which has caused over two million deaths worldwide (1). SARS-CoV-2 proteins are translated from nine canonical subgenomic mRNAs, generated by a discontinuous transcription process that results in all mRNAs having identical 5’-ends (2). To promote translation and to protect viral RNA from host surveillance by the innate immune system (3), coronaviral mRNAs are capped with guanosine monophosphate. Next, the guanosine cap is methylated by the nsp14-nsp10 heterodimer to generate Cap-0-RNA. Finally, a methyl group is transferred from *S*-adenosylmethionine (SAM) to the 2′-OH of the first adenosine residue to form Cap-1-RNA. For coronaviruses, this last reaction is catalyzed by the 2′-*O*-methyltransferase (MTase), a heterodimeric complex of nsp16 with the activator nsp10 (Fig. 1*A*) (3) (4, 5). Inhibitors of nsp16 reduce viral titer and delay the interferon response in mice, validating nsp16 as a target for the development of anti-viral small molecules (6, 7). To support drug discovery efforts, researchers around the globe have determined crystal structures of the SARS-CoV-2 nsp16-nsp10 in complex with ligands (8–12), complementing prior structural biology observations for this enzyme from SARS-CoV and MERS-CoV (13–15). This structural information has facilitated a detailed examination of the SARS-CoV-2 nsp16-nsp10 substrate binding sites (10, 11). It is well established that the 2′-*O*-MTase in SARS-CoV-2 and other RNA viruses can be activated by divalent metal ions (13, 14, 16). Yet, despite this extensive structural information, a major gap exists in understanding of the role of metal ions in the 2′-*O*-methyl transfer reaction and the position of ribonucleotides in the RNA binding groove since structures of these complexes have not been reported.

**Fig. 1.**
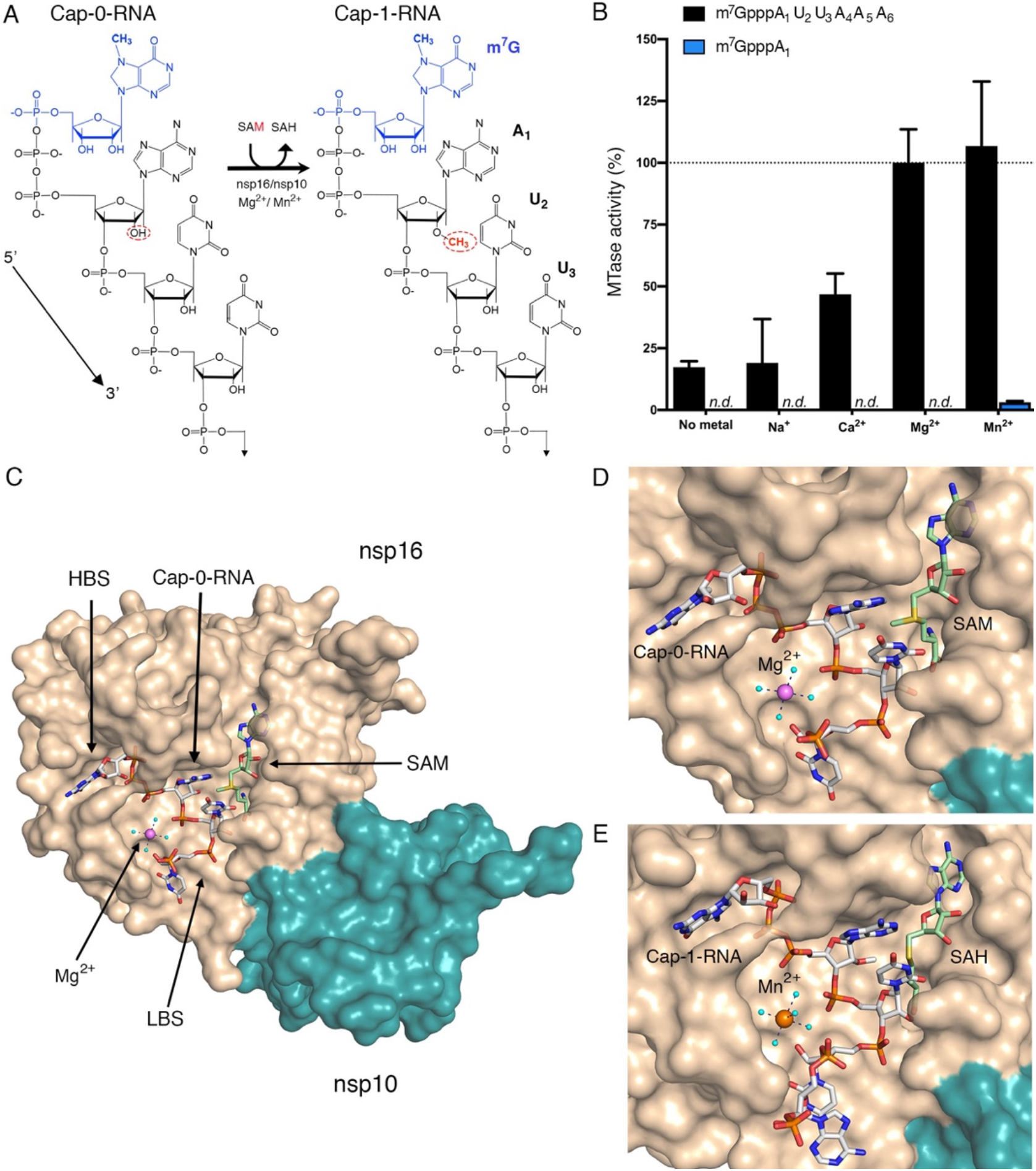
Metal ions catalyze 2′-*O*-methyl transfer and orient the Cap-RNA in the active site. *(**A**)* The schematic representation of the 2′-*O*-methyl transfer reaction. The methylated guanosine cap (m^7^G) moiety is colored in blue and the methyl group added to the A1 2′-OH position in red. The remainder of the RNA structure in black with the first three labeled ribonucleotides. *(**B**)* The MTase-Glo luminescence assay results for Cap-0-RNA (m^7^GpppAUUAAA, black bars) and Cap-0 analog (m^7^GpppA, blue bars) as substrates. *n.d.=*no activity detected. The nsp16-nsp10 activity with Cap-0-RNA, SAM and Mg^2+^ was selected as a reference (100%) and all measured activities were normalized to this value. All values are means ± the standard deviation for biological triplicates conducted in two separate experiments using two distinct preparations of nsp16-nsp10 (*n*=6). *(**C**)* The overall view of the nsp16-nsp10 in complex with Mg^2+^, Cap-0-RNA and SAM, (PDB 7JYY). The high affinity binding site is labeled as HBS and the low affinity RNA binding site as LBS. Close-up views of nsp16-nsp10 in complex *(**D**)* with Mg^2+^, Cap-0-RNA and SAM (PDB 7JYY), or *(**E**)* with Mn^2+^, Cap-1-RNA and SAH (PDB 7L6R). The nsp16 and nsp10 are represented as solvent exposed surfaces in tan and teal, respectively. Capped RNAs, SAM and SAH are shown as sticks; carbons are in grey for capped RNAs, and green for SAM and SAH, oxygens in red, nitrogen in blue, phosphates in orange, sulfur in yellow. Metal ions are shown as large spheres colored in violet for Mg^2+^ and orange for Mn^2+^. Water are small, cyan spheres. Hydrogen bonds between metal ions and waters from first hydration sphere are shown as black dashed lines.

To initiate this study, we first extended prior studies on SARS-CoV and confirmed that the SARS-CoV-2 2′-*O*-MTase also requires divalent cations. We used a custom-synthesized Cap-0-RNA substrate comprised of the *N*^7^-methylated guanosine (m^7^G) attached via a triphosphate bridge to a short RNA (AUUAAA), which matches the naturally occurring ribonucleotides at the 5’-end of SARS-CoV-2 mRNAs (17). At a concentration of 3 mM, both Mg^2+^ and Mn^2+^ significantly increased MTase activity (Fig. 1*B*). In contrast, 3 mM Ca^2+^ yielded only 50% of the activity observed with Mg^2+^ and Na^+^ did not stimulate activity. These data are consistent with observations for SARS-CoV nsp16 (4, 13, 14). The activity of SARS-CoV-2 nsp16 with the Cap-0-RNA substrate (m^7^GpppAUUAAA) is over 10-times higher than that of the Cap-0 analog (m^7^GpppA) (Fig. 1*B*, blue bars). Isothermal calorimetry (ITC) measurements were utilized to show that while it is a poor substrate for catalysis, m^7^GpppA does bind nsp16-nsp10 (K_d_ = 6.6 ± 0.3 μM) with 3-fold higher affinity than to nsp16 alone (K_d_ = 28.0 ± 5.5μM) (Table S1). In contrast, m^7^GpppG bound only to the nsp16-nsp10 complex (K_d_ = 20.0 ± 2.7 μM). We also determined the binding affinities for the methyl donor SAM and the product SAH for nsp16 and the nsp16-nsp10 heterodimer. Neither SAM nor SAH bound to nsp16 alone, but both SAM and SAH bound to the nsp16-nsp10 heterodimer with K_d_ values of 6.9 ± 1.3 μM and 13.0 ± 1.2 μM, respectively. These results from biochemical and ITC studies together, as well as work by others (4, 9, 11–13), indicated that the Cap-0 analog is capable of binding to nsp16 alone and that nsp10 greatly enhanced the binding affinity. However, a short capped RNA and the presence of Mg^2+^ or Mn^2+^ dramatically increased the rates of catalysis.

To gain further insight into how metal ions stimulate catalysis, we took a structural biology approach. Crystals of nsp16-nsp10 in complex with SAM from different crystallization conditions were soaked with the custom-synthesized m^7^GpppAUUAAA substrate in the presence of Mg^2+^ or Mn^2+^ (see methods). Multiple datasets were collected and, ultimately, three with the highest resolution and the best data statistics were selected for further analysis. Crystal #1 (PDB 7JYY) grew from a high NaCl concentration condition and was soaked with substrates and low MgCl_2_ concentration for 1.5 hours. In this crystal, we observed Cap-0-RNA, SAM, and Mg^2+^ (Fig. 1*C*, *D*). Crystal #2 (PDB 7L6R) grew from a high (NH_4_)_2_SO_4_ concentration condition and was soaked with substrates and MnCl_2_ for 6 hours. In this crystal, we observed Mn^2+^ and the products of the reaction, Cap-1-RNA and SAH, indicating that the methyl transfer reaction occurred in the crystal (Fig. 1*E*). Crystal #3 (PDB 7L6T) grew from a high magnesium formate concentration condition and was soaked with substrates for 6 hours. In this crystal we observed two Mg^2+^ ions and products of the reaction, Cap-1-RNA and SAH. The first Mg^2+^ occupied the same metal binding site as in Crystals #1 and #2 and the second Mg^2+^ directly interacted with phosphate groups of the capped RNA (Fig. S1). The number of ribonucleotides in Crystal #1 and #3 included the cap and first three ribonucleotides (m^7^GpppAUU) and the phosphate group of the fourth nucleotide. Crystal #2 contained the whole A_4_ ribonucleotide and the phosphate group of A_5_.

An overlay of the previously reported Cap-0 analog (PDB 6WRZ (10)) and Cap-0-RNA (PDB 7JYY, this study) structures showed they are very similar with a root-mean-square-deviation of 0.33 Å (Fig. 2*A*). We previously showed that the cap binding site, also called the High Affinity Binding Site (HBS), is bordered by flexible loops that adopt an open conformation upon Cap-0 analog binding (10). Interactions of the nsp16 residues Tyr6828, Tyr6930, Lys6935, Thr6970, Ser6999 and Ser7000 with the Cap-0 analog and Cap-0-RNA are similar in both structures. The m^7^GpppA in the HBS is stabilized by stacking of the m^7^G and A_1_ bases with Tyr6828 and Tyr6930 residues, respectively (Fig. 2*A*). The O2′ of the A_1_ ribose interacts with the conserved nsp16 catalytic residues (13, 14), corresponding to Lys6839-Asp6928-Lys6968-Glu7001 in our structures (Fig. 2*B*), as well as with the conserved water molecule we previously identified (10). The interactions between Asn6841, SAM and O2′ from the A_1_ are also consistent between these structures.

**Fig. 2.**
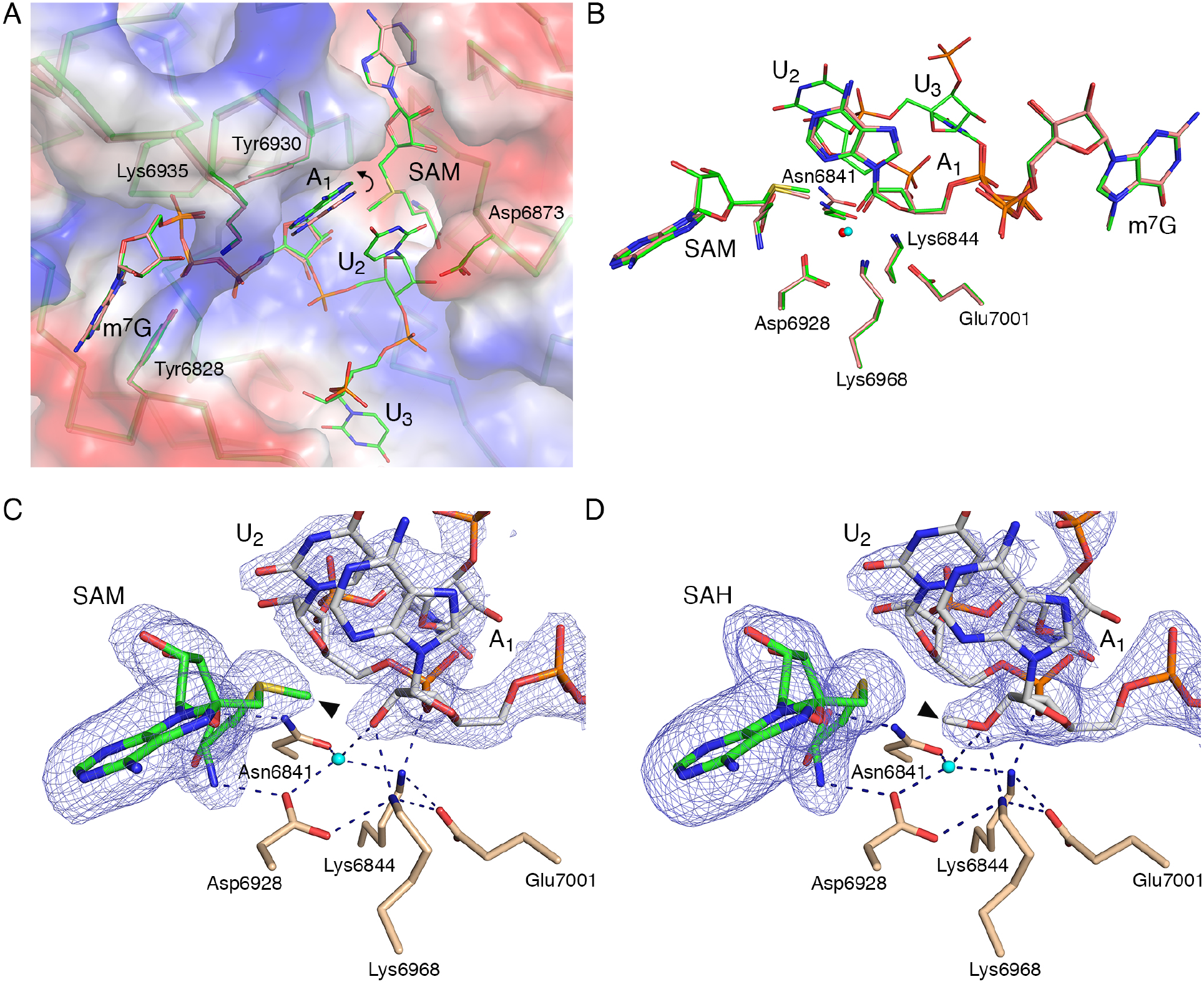
The catalytic site of the SARS-CoV-2 2′-*O*-MTase. *(**A**)* Superimposition of the Cap-0 analog (PDB 6WVN, salmon) and Cap-0-RNA (PDB 7JYY, green) structures with plotted electrostatic surface shown in blue (positive charge) and red (negative charge). Selected residues of nsp16, SAM, the Cap-0 analog and Cap-0-RNA are labeled and shown as sticks; carbons in salmon and green for Cap-0 analog and Cap-0-RNA, respectively, with oxygens in red, nitrogens in blue, and sulfurs in yellow. Repositioning of A1 base is marked with a curved arrow. *(**B**)* Catalytic residues of nsp16, Cap-0 analog, Cap-0-RNA and conserved water for the same structures and same color scheme as in panel ***A***, with waters are shown as small spheres in red and cyan for PDBs 6WVN and 7JYY, respectively. *(**C-D**)*. Wall-eyed pseudo-stereo view of the active sites for complexes of nsp16-nsp10 with Cap-0-RNA/SAM (PDB 7JYY) on the left and nsp16-nsp10 with Cap-1-RNA/SAH (PDB 7L6R) on the right. The catalytic site residues, SAM and capped RNAs are labeled and shown as stick models with atoms colored in wheat, green, and grey for carbons of nsp16, SAM and capped RNA, respectively, with red for oxygens, blue for nitrogens, yellow for sulfurs. Conserved catalytic waters are shown as cyan spheres, hydrogen bond interactions as black, dashed lines, and the omit |Fo-Fc| electron density maps contoured at the 3σ level as blue mesh. The methyl group of SAM and Cap-1-RNA are marked with black triangles.

Of particular note was the space between the Tyr6930 and Asp6873 side chains, which is occupied by the A_1_ base in the Cap-0 analog structure. In the Cap-0-RNA structure, this space accommodates the stacked bases of A_1_ and U_2_, with the U_2_ base forcing repositioning of the A_1_ ribonucleotide (Fig. 2*A*). Superposition of the Cap-0 and Cap-0-RNA structures revealed that this repositioning involves: i) a 0.6 Å shift of the A_1_ base towards the side chain of Tyr6930 without notable changes in the positions of Asn6873 and Tyr6930, and ii) a 0.4 Å decrease in the distance between O2′ of A_1_ and the SAM methyl group (Fig. 3*A*), which occurs without significant changes in position of catalytic residues (Fig. 2*B*). The movement and alignment of the A_1_ O2′ atom toward the methyl group of SAM may explain why the additional ribonucleotides increase the efficiency of the methyl transfer reaction (18–20).

**Fig. 3.**
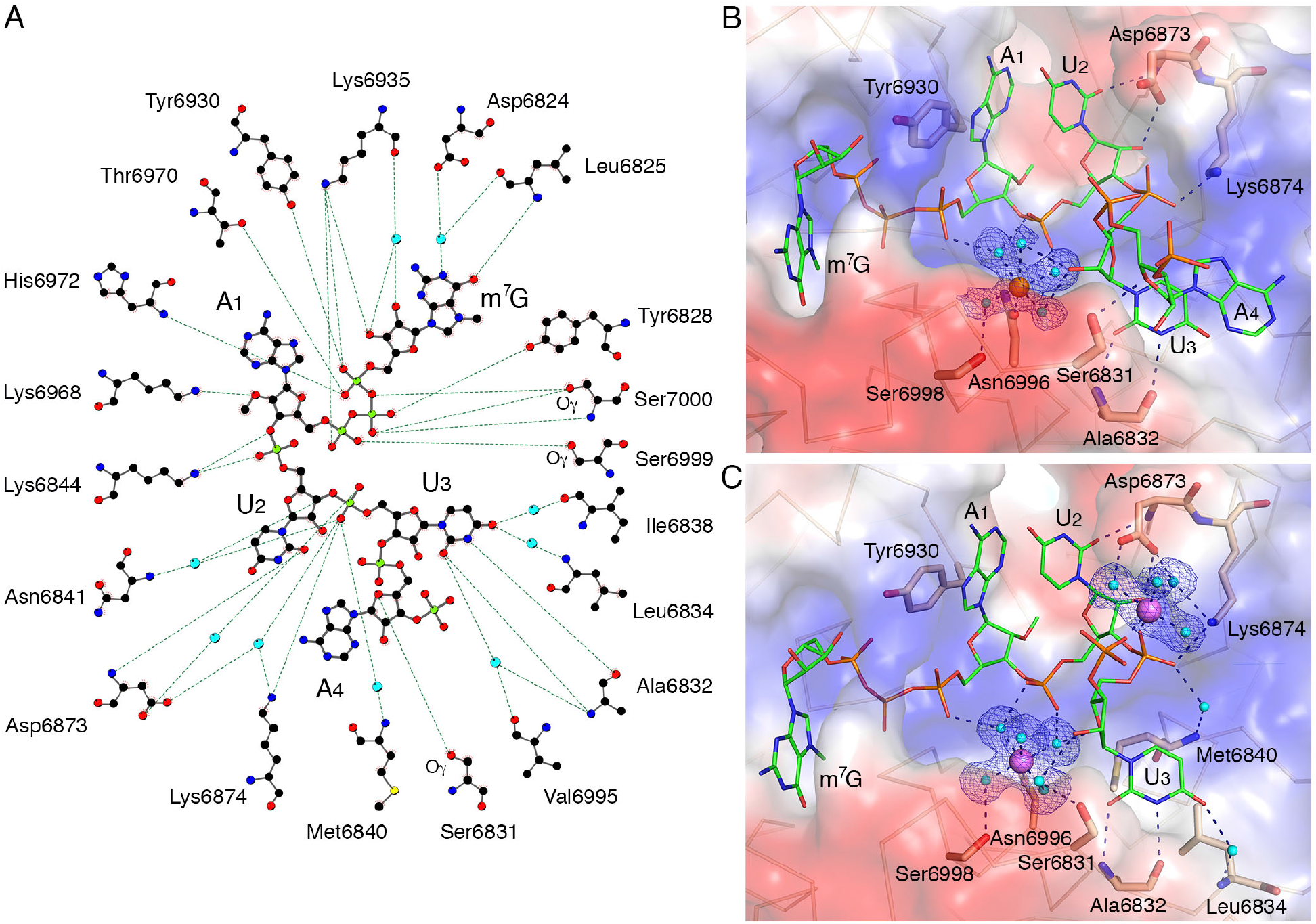
Interactions between nsp16 and Cap-1-RNA. *(**A**)* A ball-and-stick representation of the hydrogen bonding network (dashed lines) between m^7^GpppA-RNA and nsp16 residues in Crystal #2. Carbons are shown in black, nitrogens in blue, oxygens in red, phosphates in lime green, sulfurs in yellow, and waters in cyan. *(**B**)* A detailed look at the hydration sphere of Mn^2+^ (orange sphere) in Crystal #2 and *(**C**)* Mg^2+^ (purple spheres) in Crystal #3, mapped on the electrostatic surface of nsp16 (blue and red) and their interactions with water (cyan spheres) and nsp16 residues (wheat sticks). The Cap-1-RNA is represented as sticks, with carbon in green, oxygens in red, nitrogen in blue and phosphate in orange. Hydrogen bond interactions are shown as black dashed lines and the omit electron density maps as blue mesh.

Although the superposition of Cap-0 and capped RNA structures revealed differences in the position of the first adenosine, no significant deviations were observed between Cap-0-RNA and Cap-1-RNA conformations (Fig. 2*A,B*), indicating that the A_1_ base is not repositioned after the methyl transfer. The structures are essentially identical with the only difference being the methyl group, which moves from SAM to the A_1_ ribose hydroxyl group during methyl transfer (Fig. 2*C,D*).

The structures in complex with capped RNAs also revealed the importance of the low affinity binding site (LBS) residues for the conformation of the mRNA in the catalytic site. The best resolved and most complete electron density for capped RNA was observed in Crystal #2. The position of U_2_ in the active site is “locked” by multiple interactions (Fig. 3*A*). The phosphate group of U_2_ interacts with waters from the hydration sphere of the metal ion and the side chain nitrogen of Lys6844; the O2 atom of the U_2_ base interacts with the main chain nitrogen of Asp6873 and the O2′ of the U_2_ ribose makes direct interactions with one of the oxygens of the side chain of Asp6873 (Fig. 3*B*). The phosphate group of U_3_ interacts directly with the side chain nitrogen of Lys6874 and forms water mediated interactions with residues Asp6873, Lys6874, Met6840 and Asn6841. The base of U_3_ interacts directly with the main chain oxygen and nitrogen atoms of Ala6832 and forms a water bridge interaction with the nitrogen of the main chain of Leu6834. The whole nucleotide A_4_ and phosphate group of A_5_, the last ordered part of the capped RNA in the Crystal #2 structure, are solvent exposed and connected with protein via a hydrogen-bond between O4’ of the A_4_ and the side chain oxygen of Ser6831 (Fig. 3*B*). The stacking interactions between bases of U_3_ and A_4_ define the position of the A_4_ nucleotide. It is unknown if the conformation of A_4_ reflects the natural interaction of nucleotides, or if a longer mRNA would form different contacts with the nsp16-nsp10 heterodimer. However, the position of m^7^G and first three nucleotides of the capped RNA closely match in all three structures and likely represent the accurate binding mode for this part of the capped RNA.

The primary metal binding site is located near the HBS and loop1 (Fig. 1*C*) with either Mg^2+^ or Mn^2+^ occupying the same site with similar interactions (Fig. 3*B,C*). The metal ions make both direct and water-mediated interactions with sidechains of nsp16 residues and the backbone of the capped RNA. The best electron density maps were observed for Crystal #3 with two magnesium ions, both of which have near-ideal octahedral geometry (Fig. 3*C*). The Mg^2+^ in the primary metal binding site is coordinated in part by interactions with phosphate groups of the triphosphate bridge linking the cap to A_1_, the phosphate group of U_2_, and the ribose of U_3_. The second Mg^2+^ directly interacts with the U_3_ and A_4_ phosphate group oxygens and through waters with the side chain oxygens of the Asp6873 and the side chain nitrogen of Lys6874 (Fig. 3*C*). Thus, the metal ion that occupies the primary metal binding site of nsp16 properly aligns capped RNA with SAM for an efficient methyl transfer reaction. The role of metal ions in facilitating orbital alignments for efficient catalysis has been demonstrated structurally for hydride transfer reactions (19). Structural evidence for the requisite orbital alignment of substrates in RNA 2′-*O*-MTases (18) and ribozymes (20) has also been observed.

Although Mn^2+^ and Mg^2+^ are catalytic co-factors in solution (Fig. 1*B*) and were observed in our crystal structures, biochemical assay showed that Mn^2+^ best stimulated the methyl transfer reaction (Fig. 1*B*). More importantly, Mn^2+^ is present at higher concentration than Mg^2+^ in the endoplasmic reticulum and Golgi, where the SARS-CoV-2 replication vesicles are formed (21, 22). These findings suggest that Mn^2+^ may be the natural co-factor for nsp16 from SARS-CoV-2 and other coronaviruses.

All proposed *in silico* drugs that could target the residues of the LBS (9, 11–13) have relied on the crystal structures of 2′-*O*-MTases with capped RNA for DENV NS5 (PDB 5DTO (23)), VACV VP39 (PDB 1AV6 (24)) and hCMTr1 (PDB 4N48 (25)). Comparison of the RNA binding site and the capped RNA conformation from these structures with SARS-CoV-2 nsp16-nsp10 (PDB 7L6R) revealed that the conformation of the m^7^G and the location of the cap binding pockets relative to the active sites are dramatically different (Fig. S2*A-D*). In the nsp16-nsp10 and VP39 structures, the *N*^7^-methyl groups are nestled in the HBS pocket (Fig. S2*A,C*). In contrast, in NS5 and hCMTr1, for which methylation at the *N*^7^ position of the G_0_ is not required for the 2′-*O*-MTase activity (25), the *N*^7^-methyl groups are pointed toward the solvent (Fig. S2*B,D*). Although m^7^G positions do not overlap, nucleotides N_1_ and N_2_ in all these structures are closely matched, which is consistent with the conserved mechanism of action and the structure of the catalytic site (Fig. 4*A* in stereo view). In all but the nsp16-nsp10 structure, the N_2_ nucleotide is sandwiched between the N_1_ and N_3_ by base stacking interactions and these three nucleotides have limited interactions with the 2′-*O*-MTase residues of RNA binding grove. In the nsp16-nsp10 structure, all functional groups of the nucleotides N_2_ and N_3_ are involved in an integrated and complex network of hydrogen bond interactions with residues of the LBS. The U_2_ base and the ribose directly interact with main chain and side chain atoms of Asp6873. The side chain of Asp6873 occupies the space that is filled by N_3_ in all other structures. The presence of Asp6873 essentially forces the U_3_ nucleotide to move to the opposite side of the RNA binding groove, where it is involved in direct and water-mediated interactions with Ala6832 and Leu6834 (Fig. 4*A,B*). The superposition of SARS-CoV-2 nsp16-nsp10 (PDB 7JYY) with MERS-CoV (PDB 5YNM (26)) revealed almost identical conformations of the proteins and the promontory Asp6873 residue, as well as the position of the Cap moiety (Fig. 4*B*). Alignment of the amino acid sequences of nsp16 from representative coronaviruses (Fig. 4*C* and Fig. S3) showed that Asp6873 is conserved across the coronaviruses, except for feline coronavirus (F-CoV). Structural and sequence alignments of SARS-CoV-2 nsp16 with the other MTases revealed that the Asp6873 is unique to coronaviruses and located in a four-residue insertion in the loop between β1 and αA (Fig. 4*D,E*). Thus, this aspartate-containing loop and the redirection of the RNA, which is also coordinated by metals, is a unique feature that is shared across coronaviruses. Importantly, its absence in mammalian methyltransferases makes the region surrounding this residue a promising site for selective coronavirus-specific inhibitors.

**Fig. 4.**
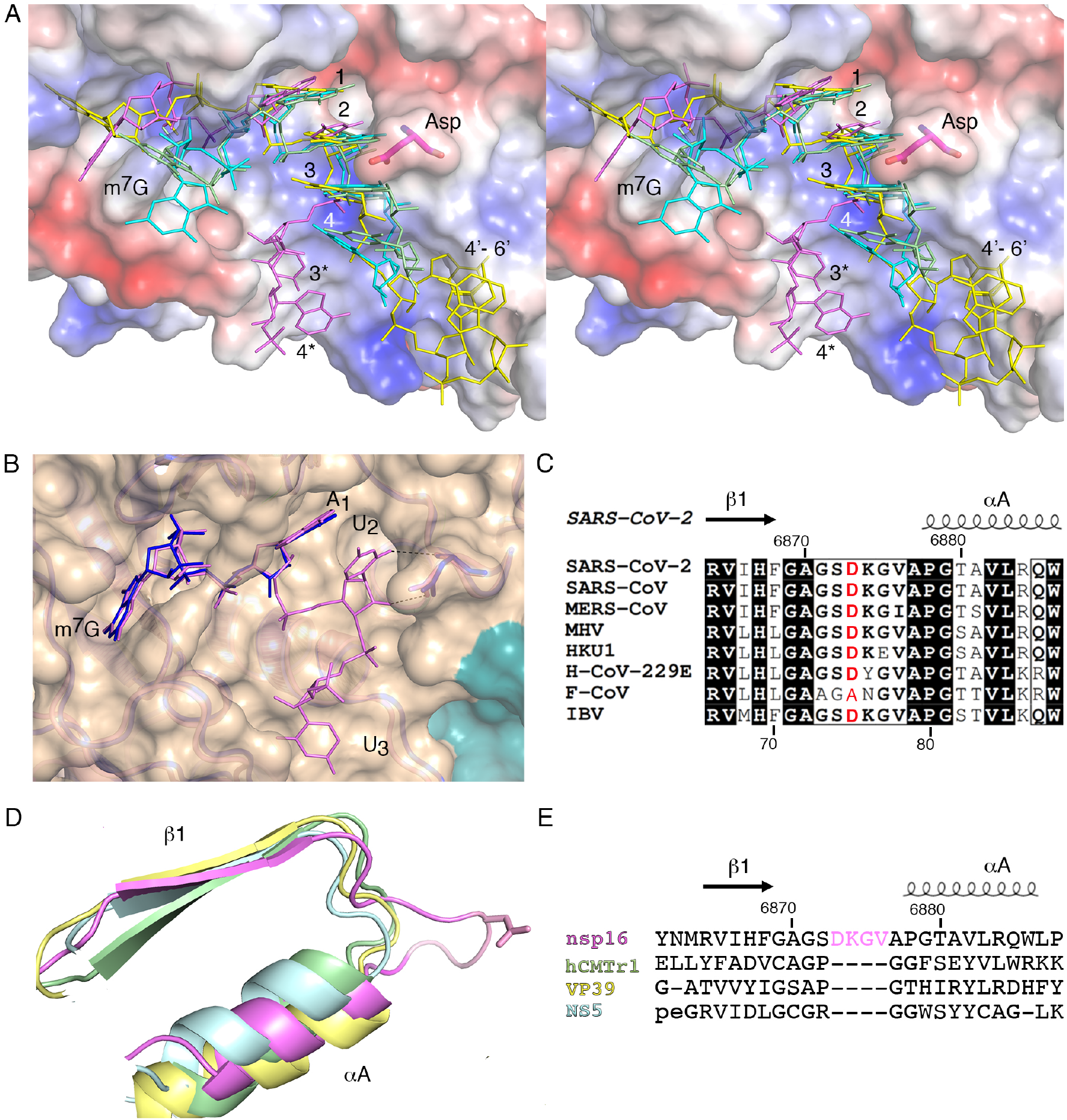
The effect of Asp6873 on conformation of the capped RNA in the nsp16 binding groove. (****E****) Wall-eyed stereo view of the superimposed capped RNAs mapped on the electrostatic potential surface of SARS-CoV-2 2′-*O*-MTase. RNAs are shown as sticks in magenta (SARS-CoV-2), cyan (DENV NS5), yellow (VACV VP39) and green (hCMTr1). The nucleotides are labeled with numbers starting from m^7^G (position 0); 3* and 4* correspond to U3 and A4 in the Cap-1-RNA from SARS-CoV-2, 4’-6’ to A4-A5-A6 in the capped mRNA from VACV. *(**B**)* Superposition of RNA binding sites of nsp16-nsp10 with Cap-0-RNA (SARS-CoV-2, magenta) and Cap-0 analog (MERS-CoV, cyan). Protein chains are shown as ribbons in orange for SARS-CoV-2 (PDB 7JYY) and green for MERS-CoV (PDB 5YNM) overlayed on the semitransparent solvent-exposed surface of nsp16 (wheat) and nsp10 (teal). Nucleotides are labeled, conserved Asp is marked with black triangle and direct hydrogen bond interactions between U2 and conserved Asp are shown as black, dashed lines. *(**C**)* Multiple sequence alignment of the loop region between I1 and IA from different coronaviruses Severe Acute respiratory Syndrome (SARS-CoV), Middle East respiratory Syndrome (MERS-CoV), Murine Hepatitis Virus (MHV), Human Coronavirus (HKU1), Human Coronavirus 229E (H-CoV-229E), Feline Coronavirus (F-CoV) and Infectious Bronchitis Virus (IBV). (***D,E***) Structural and sequence alignments of nsp16 in tan, hCMTr1 in green, VP39 in yellow and NS5 in cyan. Protein chains are shown as cartoon models with conserved Asp6873 from nsp16 shown as sticks and the insertion loop highlighted in pink.

## Materials and Methods

### Protein expression, purification, and crystallization

Recombinant nsp16 and nsp10 proteins with 6xHis-tags removed were purified from *Escherichia coli* and crystallized as previously described (10).

### MTase activity

The Cap-0 analog (m^7^GpppA) was obtained from New England Biolabs and the Cap-0-RNA (m^7^GpppAUUAAA) was custom-synthesized. The MTase activity was measured using the MTase-Glo Methyltransferase bioluminescence assay (Promega) (27) in buffer conditions suitable for use with SARS-CoV-2 nsp16. Luminescence was measured using a TECAN Safire2 microplate reader in arbitrary units and normalized assigning 100% to the activity in presence of Mg^2+^ and Cap-0-RNA. The plot was created using GraphPad Prism V9 and shows the average and the standard deviation of three measurements using two different protein purifications.

### Isothermal Titration Calorimetry (ITC)

Binding affinity was determined using a MicroCal PEAQ-ITC system (Malvern, Worcestershire, UK) at 25° C. The sample cell volume was 200 μL and the total syringe volume was 40 μL. For each titration, the first injection was performed using 0.4 μl which was then followed by 18 additional injections at 2 μl per injection. The first injection was considered a void and was automatically removed from data analysis. Each injection was spaced by 120 s after a 60 s initial delay. SARS-CoV-2 samples of nsp10, nsp16 and nsp16-nsp10 were individually loaded into the sample cell and then titrated with either SAH, SAM, m^7^GpppA Cap analog or the m^7^GpppG Cap analog (New England Biolabs). All samples were dialyzed overnight in ITC buffer (200 mM NaCl, 50 mM HEPES (pH 8.0), 0.1 mM ZnCl_2_, and 1 mM DTT). The concentrations of SARS-CoV-2 nsp10 or nsp16 used in experiments were 200 μM and 40 μM, respectively, and the concentrations of all substrates used were 500 μM. For titration experiments of the SARS-CoV-2 nsp16-nsp10, the two proteins were mixed to the final concentrations 25μM and 200 μM, respectively, and incubated for 30 mins at room temperature. A titration of SARS-CoV-2 nsp10 into nsp16 was performed at the 8:1 ratio to ensure no enthalpy was detected for the complex formation alone. The SARS-CoV-2 nsp16-nsp10 was titrated with substrates SAH and SAM at 375 μM and each Cap analog at 280 μM. Individual titration data were analyzed with MicroCal PEAQ-ITC Analysis Software using a single-site binding model and non-linear curve fitting. Each experiment was performed in triplicate and the resulting values and standard error in the fitted parameters for n, K_d_, ΔH, ΔTS, and ΔG were obtained and are summarized in Table S1.

### Crystal soaking experiments

Crystal #1 was grown from 0.1 M sodium citrate (pH 5.6) and 1.0 M ammonium dihydrogen phosphate, soaked for 1.5 h with 0.2 mM Cap-0-RNA, 5 mM SAM and 5 mM MgCl_2_ and cryoprotected with 4 M sodium formate. Crystal #2 was grown from 0.1 M citric acid (pH 5.0) and 0.8 M ammonium sulfate and was soaked for 6 hours with 0.2 mM Cap-0-RNA, 5 mM SAM and 20 mM manganese chloride. Crystal #3 was grown from 0.1 M HEPES (pH 7.5) and 0.5 M magnesium formate and was soaked for 6 hours with 0.2 mM Cap-0-RNA, 5 mM SAM. Crystals #2 and #3 were cryoprotected with 25% sucrose in respective reservoir solutions.

### Data collection, processing, structure solution and refinement

Data sets were collected at the beam lines 21ID-D and 21ID-F of the Life Sciences-Collaborative Access Team (LS-CAT) at the Advanced Photon Source (APS), Argonne National Laboratory. Images were indexed, integrated and scaled using HKL-3000 (28). Data quality and structure refinement statistics are shown in Table S2. All structures were determined by Molecular Replacement with Phaser (29) from the CCP4 Suite (30) using the crystal structure of the nsp16-nsp10 heterodimer from SARS-CoV-2 as a search model (PDB ID 6W4H). The initial solutions went through several rounds of refinement in REFMAC v. 5.8.0266 (31) and manual model corrections using Coot (32). The water molecules were generated using ARP/wARP (33) followed by an additional rounds of refinement in REFMAC. All structures were carefully examined, and three data sets were selected for further structural studies. For all structures, the Cap-0-RNA, SAM and Mg^2+^ were fit into electron density maps and further refined. Inspection of anomalous and Fourier difference electron density maps revealed that, for Crystal #1 the nsp16-nsp10 heterodimer formed the complex with the Cap-0-RNA, SAM and Mg^2+^, for Crystal #2 the complex was formed with the Cap-1-RNA, SAH and Mn^2+^, and for Crystal #3 the complex was formed with the Cap-1-RNA, SAH and two Mg^2+^. In Crystals #1 and #3, no additional electron density was detected beyond phosphate group of A_4_. In Crystal #2, the presence of well-defined electron density near phosphate group of A_4_ allowed unambiguously extending of the RNA model by adding sugar and base for A_4_ and phosphate group of A_5_. All structures were further refined with the Translation–Libration–Screw (TLS) group corrections, which were created by the TLSMD server (34). The quality control of the models during refinement and for the final validation of the structures were done using MolProbity (35) (http://molprobity.biochem.duke.edu/). All structures were deposited to Validated SARS-CoV-2 related structural models of potential drug targets (https://covid19.bioreproducibility.org/) and to the Protein Data Bank (https://www.rcsb.org/) with the assigned PDB codes 7JYY (Crystal #1), 7L6R (Crystal #2) and 7L6T (Crystal #3). A fourth structure (PDB 7JZ0) was also determined as part of this study and deposited but was not analyzed further as the metal was modeled as Na^+^, which does not catalyze the reaction, although data on this structure are included in Table S2 for reference. All models of the structures were created in PyMOL open source V 2.1 (36), diagram of interactions was created in LigPlot+ (37).

### Structural, sequence alignment and phylogenic tree

The PDB coordinates of SARS-CoV-2 nsp16 and nsp10 were analyzed using the FATCAT (38), POSA (39) and DALI (40) servers to perform structural alignments with MERS-CoV, NS5, hCMTr1 and VP39. Generated PDB files were downloaded from the servers and modeled in PyMOL open-source V 2.1. The protein sequence of 2′-*O*-MTases were obtained from the National Center for Biotechnology Information database; feline coronavirus (F-CoV, AGT52079), murine hepatitis virus (MHV, YP_009915686.1), Human coronavirus (HKU1YP_460023.1), replicase polyprotein 1ab [Human coronavirus 229E (H-CoV-229E, AGT21344.1:6464-6763), Infectious bronchitis virus (IBV, NP_066134.1:6328-6629), DENV nonstructural protein NS5 (NS5, NP_739590.2), *Homo sapiens* (hCMTr1, BAA07893.3 KIAA0082) and VACV (VP39, NC_006998.1). The multiple sequence alignment was performed using Clustal-O (https://www.ebi.ac.uk/Tools/msa/clustalo/) and merged with the coordinates of the structure with PDB code 7JYY using ESPript 3.x (41). The phylogenic tree was created using MacVector and processed in iTol (https://itol.embl.de)

## Acknowledgments

We thank Grant Wiersum, Olga Kiryukhina and Ivgeniia Dubrovska for technical assistance in protein expression, purification and crystallization. We also thank Masoud Vedadi (University of Toronto) for helpful advice on obtaining the substrate. **Funding:**This project was funded with Federal funds from NIAID/NIH/HHS (Contract HHSN272201700060C) (to A.M. and K.S) and by the Chicago Biomedical Consortium COVID Response Award CR-003 (to K.S.) P.H. was funded by NCI R01-CA142861 (to L. Laimins).

## Author contributions

G.M. led the project and solved and analyzed the structures. M.R.L. purified protein, designed and conducted biochemical assays and analyzed experimental data. L.S. designed and conducted crystallization and soaking experiments. J.S.B. collected crystallographic data. P.H. participated in preliminary biochemical experiments. C.M.D. and A.D.M. designed and conducted binding studies and analyzed data. G.M., M.R.L., L.S., N.I. and A.D.M. wrote the manuscript. K.J.F.S. oversaw all aspects of the project and edited the manuscript.

---

**Fig. S1.**
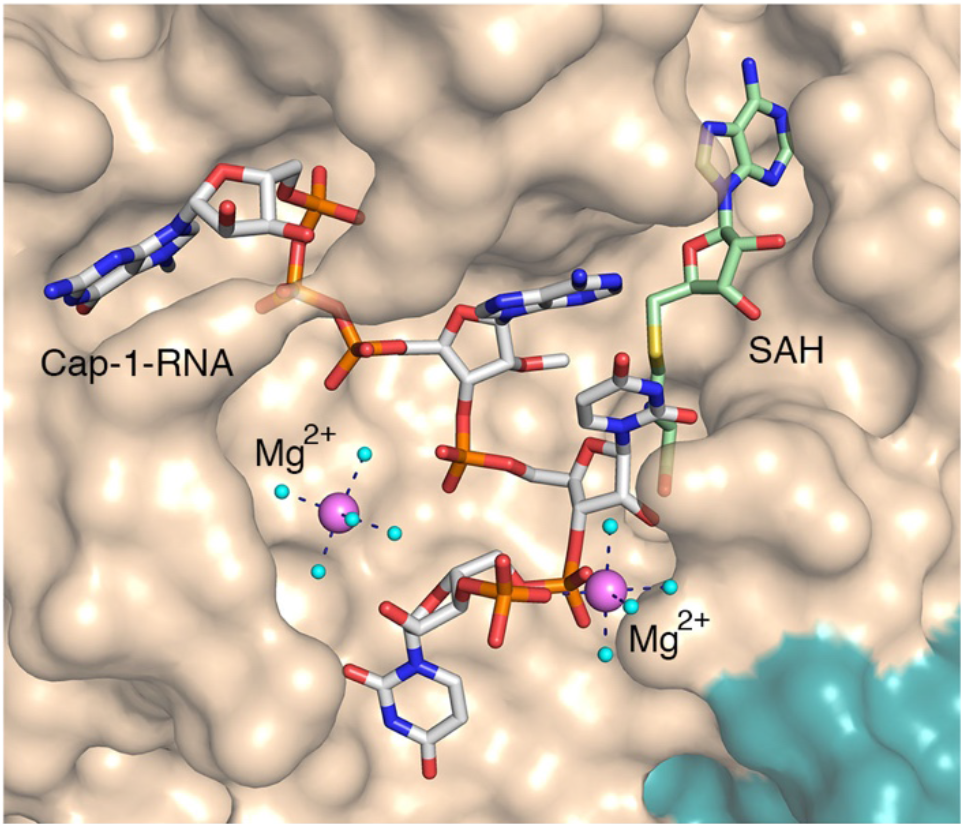
Close-up views of nsp16-nsp10 in complex with two Mg^2+^, Cap-1-RNA and SAH. The nsp16 and nsp10 (PDB code 7L6T) are represented as solvent exposed surfaces in tan and teal, respectively. Cap-1-RNA and SAH are shown as sticks; carbons are in grey for capped RNAs, and green for SAH, oxygens in red, nitrogen in blue, phosphates in orange, sulfur in yellow. Mg^2+^ are shown as large spheres colored in violet. Water are small spheres in cyan. Hydrogen bonds between metal ions and waters from first hydration sphere are shown as black dashed lines.

**Fig. S2.**
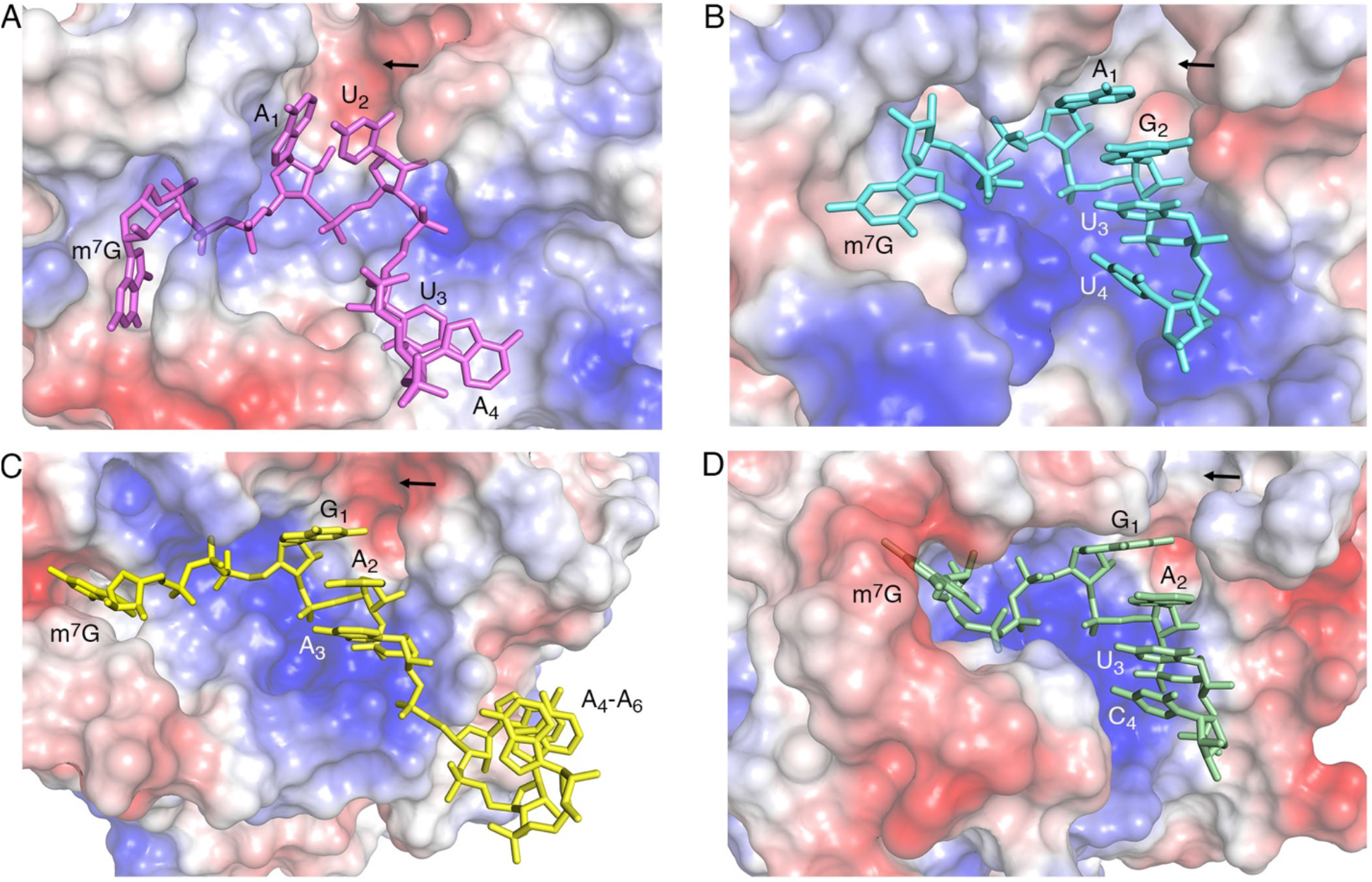
Comparison of the RNA binding grooves of 2′-*O*-MTases. Electrostatic potential surface representations of the 2′-*O*-MTases of SARS-CoV-2 nsp16-nsp10 (PDB code 7L6R) (****A****), DENV NS5 (PDB code 5DTO) (****B****), VACV VP39 (PDB code 1AV6) (****C****), and hCMTr1 (PDB code 4N48) (****D****) with the bound capped RNAs. Arrows indicate the SAM/SAH binding clefts and nucleotides are labeled as A for adenosine, G for guanosine, C for cytosine and U for uridine. The RNA bases are numbered starting from the 5’-guanine cap (m^7^G) (position 0). RNAs are shown as sticks in magenta (SARS-CoV-2), cyan (DENV NS5), yellow (VACV VP39) and green (hCMTr1).

**Fig. S3.**
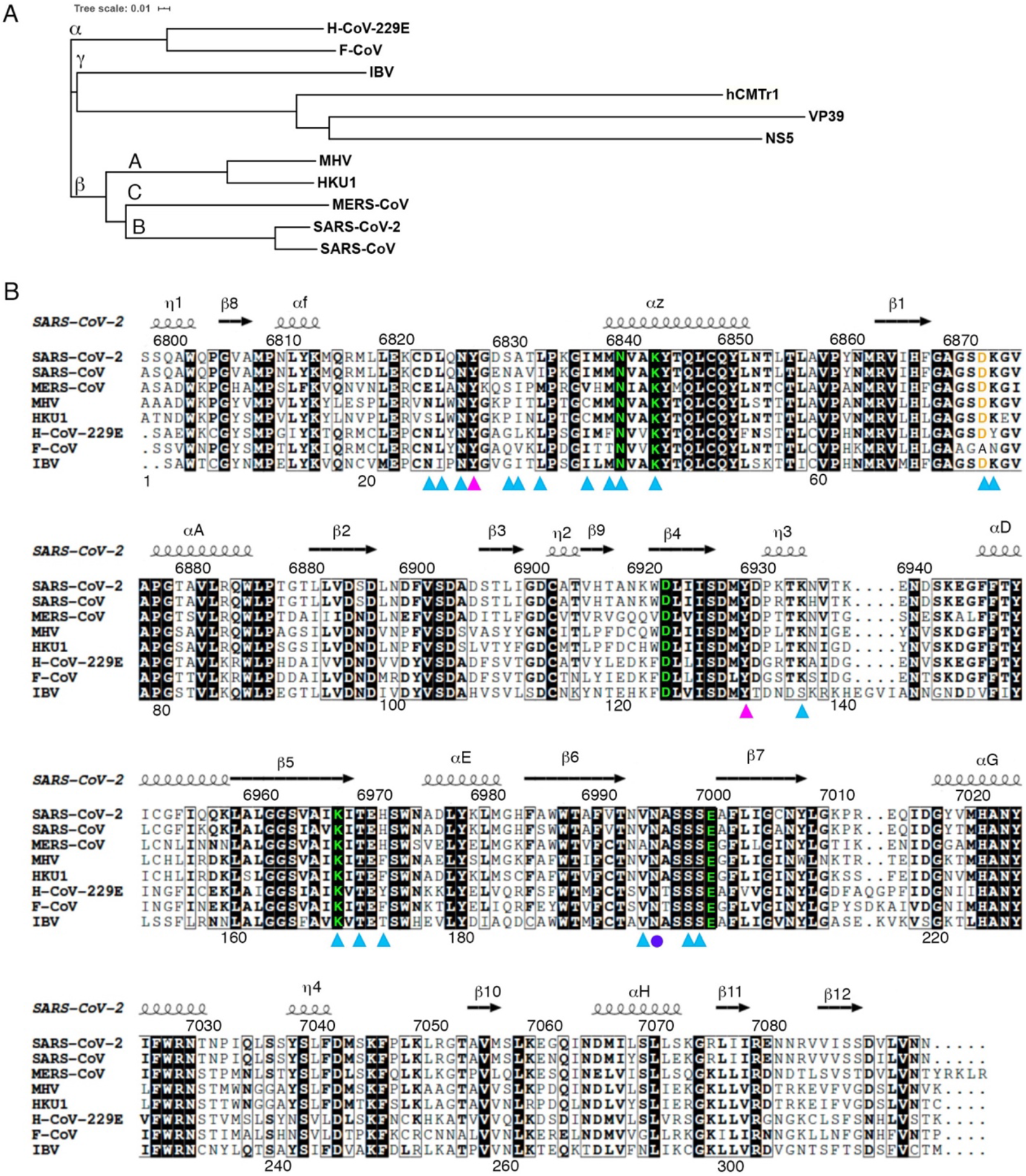
Conservation of 2′-*O*-Methyl transferases. ***(A)*** Phylogenetic tree of MTases from coronaviruses, VACV (VP3), DENV (NS5), and human (hCMTr1). ****(B)**** Primary amino acid sequence alignment of the 2′-*O*-MTases from different clades of coronaviruses showing the structural elements of nsp16 from SARS-CoV-2. Residues that are 100% conserved are shaded in black, catalytic residues are highlighted in green, residues that participate in Cap stacking are marked by pink triangles, RNA-protein interactions are marked by blue triangles, and the direct protein-metal interaction is indicated with purple circle. The conserved Asp6873 is colored orange. Abbreviations are used: Mice Hepatitis Virus (MHV), Human Coronavirus-229E (H-CoV-229E), Feline Coronavirus (F-CoV) and Infectious Bronchitis Virus (IBV).

**Table S1.**
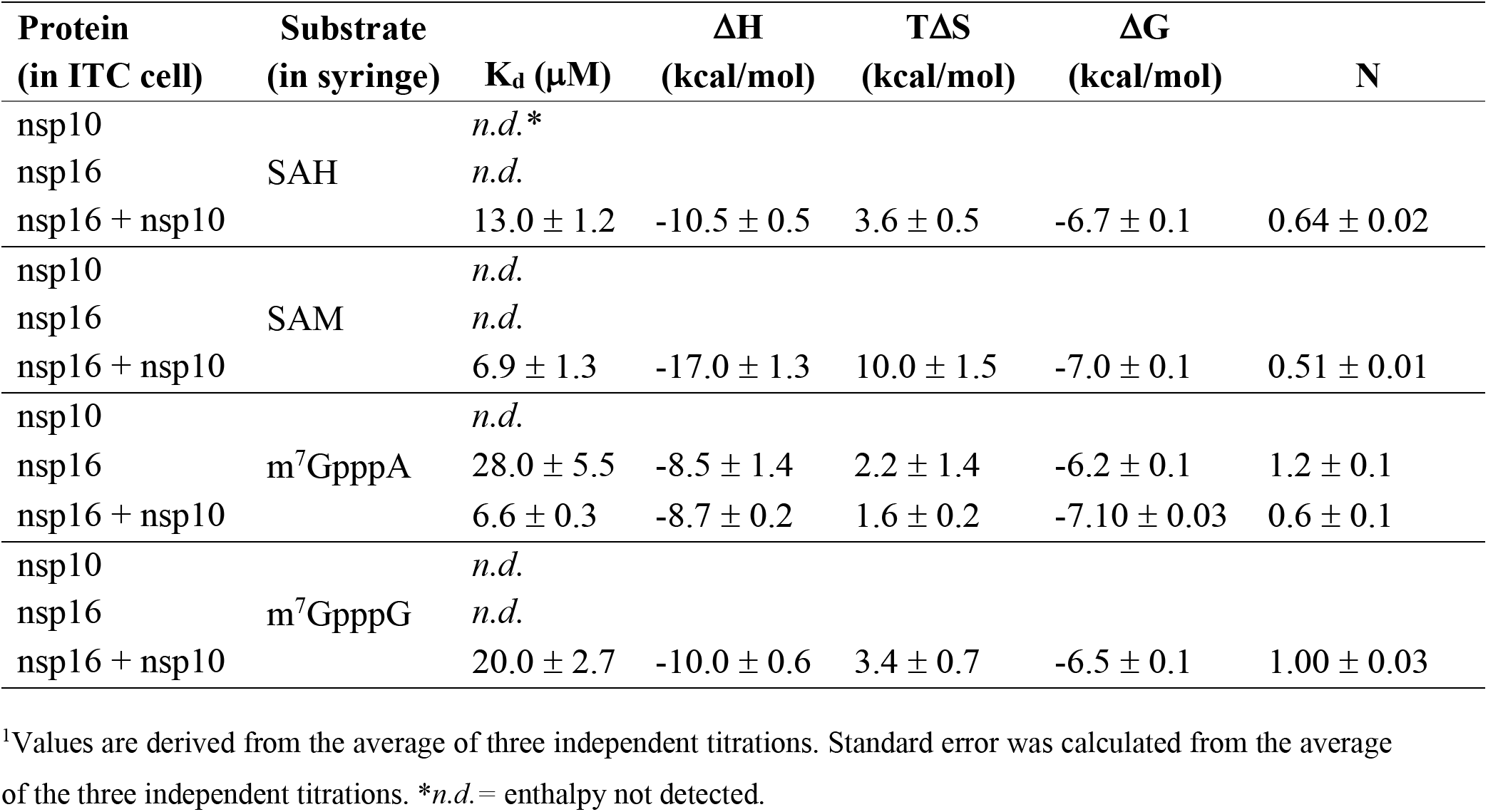
ITC derived thermodynamic parameters for the interactions of substrates with SARS-CoV-2 nsp10, nsp16 and nsp16-nsp10.

**Table S2.** Data quality and structure refinement statistics.

| **Crystal** | **#1** | **#2** | **#3** | **#4** |
| --- | --- | --- | --- | --- |
| **PDB Accession Code** | **7JYY** | **7L6R** | **7L6T** | **7JZ0** |
| **Data Collection** |  |  |  |  |
| Wavelength (Å) | 0.97872 | 1.54980 | 1.12708 | 0.97872 |
| Space group | *P3<sub>2</sub>21* | *P3<sub>1</sub>21* | *P3<sub>1</sub>21* | *P3<sub>2</sub>21* |
| Unit cell parameters (Å; °) | *a*=*b*=166.95,<br>*c*=98.81;<br>$\alpha$=$\beta$=90.00,<br>$\gamma$=120.00 | *a*=*b*=168.76,<br>*c*=52.27;<br>$\alpha$=$\beta$=90.00,<br>$\gamma$=120.00 | *a*=*b*=169.35,<br>*c*=52.63;<br>$\alpha$=$\beta$=90.00,<br>$\gamma$=120.00 | *a*=*b*=167.24,<br>*c*=98.87;<br>$\alpha$=$\beta$=90.00,<br>$\gamma$=120.00 |
| Resolution range (Å) | 30.00-2.05 (2.09-2.05) | 30.00-1.98 (2.01-1.98) | 30.00-1.78 (1.81-1.78) | 30.00-2.15 (2.19-2.15) |
| No. of reflections | 99,216 (4,938) | 59,455 (2,968) | 81,048 (3,493) | 86,204 (4,265) |
| *R*<sub>merge</sub> (%) | 8.3 (107.9) | 9.5 (120.3) | 9.0 (115.6) | 9.0 (123.9) |
| Completeness (%) | 100.0 (99.8) | 100.0 (100.0) | 96.6 (84.2) | 100.0 (100.0) |
| Rp.i.m.<sup>a</sup> | 0.032 (0.415) | 0.032 0.411) | 0.036 (0.513) | 0.036 (0.493) |
| CC<sub>1/2</sub><sup>b</sup> | 0.996 (0.765) | 0.997 (0.663) | 0.996 (0.560) | 0.994 (0.735) |
| $\langle I/\sigma(I) \rangle$ | 24.1 (2.1) | 31.2 (2.0) | 20.8 (2.1) | 26.6 (1.8) |
| Multiplicity | 7.6 (7.6) | 9.8 (9.3) | 6.8 (5.8) | 7.1 (7.1) |
| Wilson *B* factor (Å<sup>2</sup>) | 35.8 | 33.0 | 24.4 | 42.2 |
| **Refinement** |  |  |  |  |
| Resolution range (Å) | 29.97-2.05 (2.10-2.05) | 29.95-1.98 (2.03-1.98) | 29.33-1.78 (1.83-1.78) | 30.00-2.15 (2.21-2.15) |
| Completeness (%) | 99.9 (99.3) | 99.9 (99.2) | 96.8 (90.8) | 99.8 (99.1) |
| No. of reflections | 94,164 (7,228) | 56,334 (4,345) | 76,229 (5,509) | 81,779 (6,240) |
| *R*<sub>work</sub>/*R*<sub>free</sub>, (%) | 16.6/18.5<br>(24.7/26.9) | 15.0/16.6<br>(27.8/25.8) | 14.1/16.1<br>(23.9/25.6) | 17.3/19.8<br>(27.1/28.4) |
| Protein chains/atoms | 4/6,692 | 2/3,302 | 2/3,359 | 4/6,628 |
| Ligand/Solvent atoms | 275/520 | 213/341 | 166/508 | 279/393 |
| Mean temperature factor (Å<sup>2</sup>) | 47.3 | 44.0 | 33.0 | 55.9 |
| **Coordinate Deviations** |  |  |  |  |
| R.m.s.d. bonds (Å) | 0.004 | 0.005 | 0.006 | 0.004 |
| R.m.s.d. angles (°) | 1.290 | 1.272 | 1.330 | 1.194 |
| **Ramachandran plot<sup>c</sup>** |  |  |  |  |
| Favored (%) | 96.0 | 98.0 | 97.0 | 96.0 |
| Allowed (%) | 4.0 | 2.0 | 3.0 | 4.0 |
| Outside allowed (%) | 0.0 | 0.0 | 0.0 | 0.0 |

Notes: Values in parentheses are for the outer shell. <sup>a</sup>Estimated Rp.i.m. as defined by Weiss (42). <sup>b</sup>Pearson's correlation coefficient as defined by Karplus and Diederichs (43). <sup>c</sup>Validation was done using MolProbity (35).

## Notes

### Competing Interest Statement

K.J.F.S. has a significant financial interest in Situ Biosciences, LLC, a contract research organization that conducts antimicrobial testing for industrial products, including antiviral testing. This work has no overlap with the interests of the company. K.J.F.S. is a consultant for a healthcare firm on public health topics related to COVID-19 that are unrelated to this article. A.D.M. has pending intellectual property related to coronaviruses including SARS-CoV-2 but none are related to this project. A.D.M. serves as a consultant for pharmaceutical companies on topics related to SARS-CoV-2 and COVID-19 that are unrelated to this article. All other authors declare no conflicts of interest.

